# An evaluation of the interpretability and predictive performance of the BayesR model for genomic prediction

**DOI:** 10.1101/2020.10.23.351700

**Authors:** Fanny Mollandin, Andrea Rau, Pascal Croiseau

**Affiliations:** Université Paris-Saclay, INRAE, AgroParisTech, GABI; BioEcoAgro Joint Research Unit, INRAE, Université de Liège, Université de Lille, Université de Picardie Jules Verne

**Keywords:** genomic prediction, QTL mapping, Bayesian model

## Abstract

Technological advances and decreasing costs have led to the rise of increasingly dense genotyping data, making feasible the identification of potential causal markers. Custom genotyping chips, which combine medium-density genotypes with a custom genotype panel, can capitalize on these candidates to potentially yield improved accuracy and interpretability in genomic prediction. A particularly promising model to this end is BayesR, which divides markers into four effect size classes. BayesR has been shown to yield accurate predictions and promise for quantitative trait loci (QTL) mapping in real data applications, but an extensive benchmarking in simulated data is currently lacking. Based on a set of real genotypes, we generated simulated data under a variety of genetic architectures, phenotype heritabilities, and we evaluated the impact of excluding or including causal markers among the genotypes. We define several statistical criteria for QTL mapping, including several based on sliding windows to account for linkage disequilibrium. We compare and contrast these statistics and their ability to accurately prioritize known causal markers. Overall, we confirm the strong predictive performance for BayesR in moderately to highly heritable traits, particularly for 50k custom data. In cases of low heritability or weak linkage disequilibrium with the causal marker in 50k genotypes, QTL mapping is a challenge, regardless of the criterion used. BayesR is a promising approach to simultaneously obtain accurate predictions and interpretable classifications of SNPs into effect size classes. We illustrated the performance of BayesR in a variety of simulation scenarios, and compared the advantages and limitations of each.

## INTRODUCTION

The primary objective of genomic prediction is to use genomic variation, usually single nucleotide polymorphisms (SNPs), to predict phenotypes, i.e. an observable trait of an individual. In particular, genomic prediction models are widely used as an evaluation tool for genomic selection in animal breeding (1), and for the calculation of polygenic risk scores for human diseases (2). As genotyping costs have declined (3), there has been a corresponding increase in the amount of genotyping data available for analysis. In addition, lower costs and better data storage capacity have allowed for increasingly dense genotypes, up to and including whole genome sequences (WGS), which in turn have enabled sequence-level genotypes to be imputed for individuals genotyped using lower density chips(4). However, analyzing these increasingly large genotype data can come at a high computational cost and requires suitable statistical methods. Although the use of higher density genotypes was initially thought to hold promise for improved prediction accuracy, their performance was not found to improve that of high density chips in real data, due to the inclusion of a large number of non-causative SNPs (5). While the exhaustive use of WGS variants has not led to meaningful improvements in prediction, they do allow for the direct inclusion of candidate, or even causal, mutations (6). For simplicity, we refer to such mutations as quantitative trait loci (QTL) throughout. If such QTLs are known a priori or can be directly identified through variable selection in the model itself, this could potentially lead to the double advantage of improving both the accuracy and interpretability of genomic prediction models(7; 8). With this in mind, custom chips, which include SNPs from a medium-density chip (intended to cover the genome) as well as candidates or causal mutations for a set of traits, have been developed, offering the cost and computational advantages of a reasonably sized chip with the increased predictive ability and interpretability provided by the inclusion of potential causal mutations.

Most models used in routine genomic selection are based on linear models, notably best linear unbiased prediction (BLUP) and genomic BLUP (GBLUP). These models assume that all SNPs contribute equally to the genomic variance, with each SNP effect following a normal distribution with common variance. Although the assumption about common SNP effects allows for great computational efficiency, it is quite strong and can limit the biological interpretability of results. To address this limitation, although deep learning models have recently started to appear (9; 10), a more frequent alternative is the set of non-linear Bayesian models comprising the so-called Bayesian alphabet. These include, among others, BayesA (1), BayesB (1), BayesC*π* (11), BayesR (12), and BayesRC (13). The aim of all of these models is to improve predictive accuracy by better estimating SNP effects through more flexible prior specifications. For instance, in the earliest model introduced, BayesA, all markers are assumed to be drawn from a normal distribution whose variance follows an Inv-*χ^2^* distribution. Although the assumptions of BayesA are arguably closer to reality than BLUP or GBLUP, it is computationally expensive to estimate variances for every SNP in dense genotyping data. Instead, a useful alternative is to assume that a (potentially large) portion of markers contribute no genetic variance. This is the strategy employed by both BayesB and BayesC, which model marker effect variances as a zero-inflated distribution by assigning null effects with a fixed probability, and assuming the variance of non-null SNPs respectively follow a per-SNP or common Inv-*χ*^2^ distribution. BayesC*π* further assumes that the proportion of null SNP effects is itself a random variable, and otherwise uses a common prior distribution for non-null SNP effects. BayesR provides additional flexibility by defining four classes of SNP effect size (null, small, medium, large), where SNP effects are modeled using a four-component normal mixture model. The related BayesRC model further allows for SNPs to be grouped into disjoint categories (e.g., according to prior biological information), for which the BayesR model is subsequently fit independently.

Although these Bayesian genomic prediction models are mainly used for phenotype prediction, they also provide valuable per-SNP information, including posterior estimates of effect size and variance, which could be used for QTL mapping. In contrast to genome-wide association study (GWAS) methods, SNP effects are estimated simultaneously and make use of variable selection within the model itself, rather than relying on univariate hypothesis tests and corrections for multiple testing. As the quantity and quality of prior biological knowledge continues to improve and the identification of causal mutations from WGS data (14) becomes increasingly feasible, the flexible model definition of BayesR and BayesRC thus make them interesting candidates for simultaneously providing good predictability and biologically interpretable QTL mapping results. In this spirit, Moser *et al*, showed encouraging results for the use of BayesR in complex traits for prediction and QTL mapping in real data (15). However, a comprehensive simulation study investigating the interpretability and performance of BayesR in a wide variety of settings is currently lacking in the literature. In addition, to date there has been little discussion of the various criteria that can potentially be used to map QTLs using the BayesR model output.

To address this gap, our goal in this work is to identify the coherence between the BayesR model specification and known QTL effects in simulated data under a variety of conditions. The BayesR approach is of particular interest here, as it has been shown in the literature to improve prediction accuracy (16), but its ability to correctly assign QTLs to the appropriate effect size categories has not yet been extensively evaluated in simulations. We focus on the case where a prior categorization of markers (i.e., the BayesRC approach) is not available. Using simulated data, we evaluate the robustness of BayesR under a wide variety of genetic architectures, phenotype heritabilities, and polygenic variances, and we illustrate the conditions under which BayesR successfully identifies known QTLs while maintaining high accuracy for phenotypic prediction. Finally, we describe and compare several statistical criteria that can be used to perform QTL mapping using BayesR output. Based on the results of our simulation study, we discuss the optimal framework for an accurate and interpretable analysis using BayesR, as well as its limitations.

## MATERIALS AND METHODS

### Data simulation based on real genotypes

To maintain a realistic linkage disequilibrium (LD) structure among SNPs, we generated simulated data based on a set of genotypes assayed using Illumina Bovine SNP50 BeadChip arrays from *n* = 2605 Montbéliarde bulls. We divided individuals into learning and validation sets (i.e., the “holdout method”), with the 80% oldest bulls (*n*_learning_ = 2083) in the former and the 20% youngest (*n*_validation_ = 522) in the latter to reflect the strategy typically used in routine genomic selection. We excluded SNPs with a minor allele frequency (MAF) less than 0.01, leaving a total of *p* =46,178 SNPs.

To simulate phenotypes **y** for the *n* = 2605 bulls, we made use of a standard linear model:

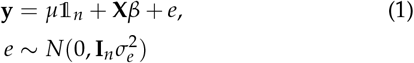

where *μ* denotes the trait mean (including fixed effects), *β* the vector of effects for the *p* SNPs, **X** the centered and scaled genotype design matrix, and *e* the residuals, assumed to follow a normal distribution with variance 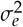. Parameters for this linear model were set as follows. For each simulated dataset we sampled from the available SNPs a set of nQTL QTLs and a set of *n*_poly_ polygenic SNPs, as well as their corresponding genetic variances for each selected marker. To reduce the impact of extreme MAFs on genomic prediction (17) and QTL detection, we focused on frequent QTLs by drawing the *n*_QTL_ and *n*_poly_ SNPs from those with a MAF ≥ 0.15. In all simulations, we selected a total of *n*_QTL_ = 5 large QTLs, varying the corresponding proportion *k* of total genetic additive variance 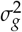 as described below. The phenotypic variance and mean were respectively set to 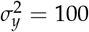 and *μ* = 0, and SNP heritability 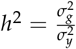 was varied across simulation settings.

**Table 1.** Simulation settings for each of the 13 QTL effect-size scenarios considered for each given level of heritability, *h*^2^ = {0.1,0.3,0.5,0.8}. The number of simulated QTLs, number of polygenic SNPs, percentage of genetic variance attributed to each QTL, and percentage of genetic variance attributed to each polygenic SNP are provided. Summing the percentage of genetic variance explained by the total number of QTLs and polygenic SNPs yields 100%.

|  |  |  |  |  |  |  |  |  |  |  |  |  |  |
| --- | --- | --- | --- | --- | --- | --- | --- | --- | --- | --- | --- | --- | --- |
| Number of QTLs | 5 | 5 | 5 | 5 | 5 | 5 | 5 | 5 | 5 | 5 | 5 | 5 | 5 |
| Number of polygenic SNPs | 9637 | 9550 | 9500 | 9450 | 9350 | 9250 | 9100 | 9000 | 8750 | 8500 | 8250 | 8000 | 7500 |
| Per-QTL % of $\sigma_g^2$ | 0.725 | 0.9 | 0.10 | 0.11 | 0.13 | 0.15 | 0.18 | 2 | 2.5 | 3 | 3.5 | 4 | 5 |
| Per-polygenic SNP % of $\sigma_g^2$ | 0.01 | 0.01 | 0.01 | 0.01 | 0.01 | 0.01 | 0.01 | 0.01 | 0.01 | 0.01 | 0.01 | 0.01 | 0.01 |

We constructed 13 scenarios with different proportions *k* of genetic variance attributed to the QTLs, with 10 independent datasets created for each (Table 1). For the SNPs randomly selected as QTLs and polygenics SNPs, the corresponding effect *β_i_* for selected SNP *i* was set as follows:

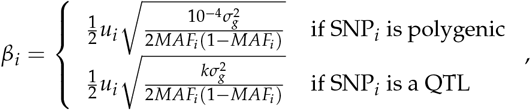

where *u_i_* was drawn from a discrete Uniform {–1,1} distribution to allow non-null effects to take on positive or negative values. For unselected SNPs (i.e., null SNPs), *β_i_* was set to 0. We varied the proportion of genetic variance attributed to each QTL between *k* =0.725% and 5%, with a greater density of values evaluated between 0.725% and 2%; we focused in particular on this range as it corresponds to more plausible QTL sizes and facilitated a study of the sensitivity of BayesR to small changes. For each value of *k*, the same *n*_QTL_ = 5 QTLs were used across scenarios, but the number (and thus the subset) of polygenic SNPs used varied (see Table 1). As the same 5 QTLs were simulated across scenarios for each of the 10 independent datasets, a total of 50 QTLs was considered. Finally, each scenario was run for four different levels of heritability *h*^2^ = {0.1,0.3,0.5,0.8}, and we evaluated the performance of BayesR for two alternatives: (1) using genotype data that excludes the 5 known QTLs, resembling a classic 50k genotyping array (“50k data”); and (2) using genotype data that includes the 5 known QTLs, which mimics a custom 50k genotyping array (“50k custom data”). In total, this corresponds to 13 × 10 × 4 × 2 = 1040 simulated datasets.

### Statistical Analysis

#### BayesR genomic prediction model

The models of the Bayesian alphabet are all based on the linear model in Equation (1). BayesR assumes that SNP effects *β_i_* follow a four-component normal mixture, making it well-aligned to our simulations (for which SNPs fall into null, weak, and strong classes). The effect of SNP *i* is assumed to be distributed as

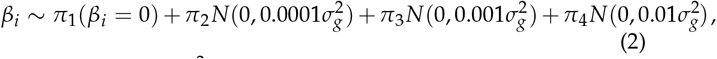

where as before, 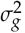 represents the total additive genetic variance (i.e., the cumulative variance of all SNP effects) and *π* = (*π*_1_, *π*_2_, *π*_3_, *π*_4_) the mixing proportions such that 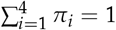. The mixing proportions *π* are assumed to follow a Dirichlet prior, *π* ~ Dirichlet(*α* + *γ*), with *α* representing a vector of pseudocounts and *γ* the cardinality of each component. In this work, we used a flat Dirchlet distribution, with *α* = (1,1,1,1), for the prior. As suggested by Moser *et al*. (15), 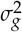 is assumed to be a random variable following an Inv—*χ*^2^ distribution.

As exact computation of the posterior distribution is intractable for this model, Bayesian inference is performed by obtaining draws of the posterior using a Gibbs sampler; full details of the algorithm can be found in (15) and (18). In practice, at each iteration of the algorithm, SNPs are assigned to one of the four categories, and their effect is subsequently sampled from the full conditional posterior distribution for the corresponding mixture component. Model parameters are then estimated using the posterior mean across iterations, after excluding the burn-in phase and thinning draws. Here, the Gibbs sampler was run for a total of 50,000 iterations, including 20,000 as a burn-in and a thinning rate of 10.

In this work, we used the open source Fortran 90 code described in (15) and available at https://github.com/syntheke/bayesR. We made a few modifications to this code, notably adding the posterior variance of estimated SNP effects at each iteration to the output; our modified BayesR code may be found at https://github.com/fmollandin/BayesR_Simulations.

Prediction accuracy for BayesR was quantified using the Pearson correlation between the true phenotypic values (**y**) and those estimated using BayesR 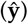 in the validation set.

#### Statistical criteria for QTL mapping

In this section, we present several potential criteria based on BayesR output that can be used for the purpose of QTL mapping. We have sub-divided these criteria into those defined for (1) each SNP individually; (2) neighborhoods, or sliding windows, around each marker; and (3) those used for ranking potential QTLs.

##### Mapping criteria for individual SNPs

BayesR is unique in the Bayesian alphabet, in that it assigns SNPs to one of four effect size classes at each iteration by weighting according to their likelihood of belonging to each. We thus have access to the posterior frequency with which SNPs were assigned to each class, which can be interpreted as an inclusion probability. We denote the posterior inclusion probability (PIP) of SNP *i* belonging to class *j* as 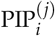, such that 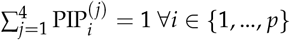. In the following we interchangeably refer to the null, small, medium, and large classes as *j* = 1, 2, 3, and 4, respectively. The PIP provides a straightforward method for classifying SNPs as having a null, small, medium, or large effect. We define the maximum a posteriori (MAP) rule for SNP *i* as

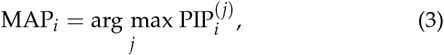

implying that SNPs are assigned to their most frequently assigned class. Since SNPs may move frequently from one class to another, the MAP in Equation (3) may not detect SNPs that are predominantly included in the model but move between the three non-null classes. Merging the non-null classes addresses this problem, and leads to a less stringent criterion, the non-null MAP:

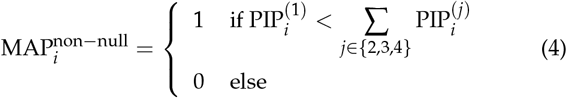

Based on this criterion, SNP *i* is thus included in the model if 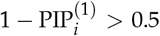. In this way, all SNPs preferentially assigned to the null class take on a value of 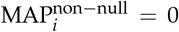, while those assigned to any non-null class (small, medium, or large) take on a value of 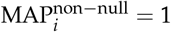.

The BayesR model definition explicitly allows for some SNPs to have larger estimated variances than methods such as GBLUP, which tends to shrink the variance of causal marks due to the assumption of a common variance (18). As such, BayesR has the potential for more closely approximating the true variance of QTLs. The posterior variance of SNP *i* corresponds to

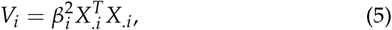

where *X_.i_* represents the *i*^th^ column of the centered and scaled genotype design matrix. As the SNP effects are computed on the scaled and centered genotype design matrix *X*, the per-SNP posterior variance can be estimated using

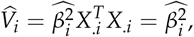

where 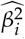 corresponds to the posterior mean of 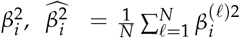, where *n* is the number of iterations and 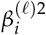 the value of 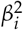 at iteration *ℓ*. We indirectly estimated this parameter as the sum of the posterior variance and squared posterior mean of each per-SNP effect. We can then estimate a posteriori the proportion of genetic variance of a SNP *i* as 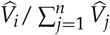.

##### Neighborhood-based mapping criteria

LD represents a preferential association between two alleles and can have a large impact on how estimated variances are distributed among SNPs in an LD block. This in turn affects the evaluation of the variance in the neighborhood of a causal mutation, as well as the ability to perform QTL mapping using the aforementioned criteria, for several reasons. First, SNPs in close proximity to a QTL are likely to be in high LD with it, and thus may erroneously have their own effects overestimated to the detriment of the QTL’s. The per-SNP criteria defined above risk incorrectly identifying a QTL as null in such cases. An alternative approach is to define a neighborhood-based criteria around each marker, thus mapping QTLs when one or more or its close neighbors is detected. Here, we define each neighborhood as a sliding window of 15 SNPs (covering approximately 1Mb) centered around each marker.

Using these neighborhoods, we define the vector of PIPs for a neighborhood centered on SNP i as follows:

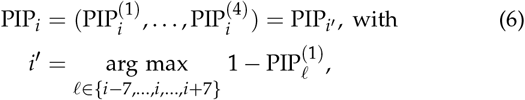

with the corresponding neighborhood inclusion probability equal to

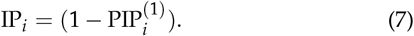

The criteria proposed in Equations (3)–(5) can thus be adapted to accommodate neighborhoods as follows:

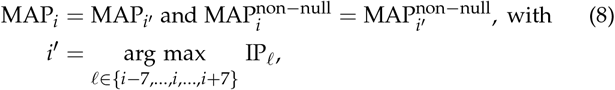

where SNP indices are assumed to be ordered according to their physical location. Similarly, the estimated variance of a neighborhood is fixed to the maximal value of its individual markers:

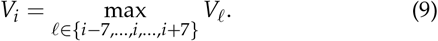

LD structure raises an additional related problem - in some cases, the BayesR algorithm may alternate assigning different SNPs in an LD block to the large effect class, which has the consequence of diluting variance over a region rather than for a single marker. The window-based criteria in Equations (8)–(9) successfully flag regions where a single SNP sufficiently stands out, but not necessarily those including several diluted effects. In addition, it can be difficult to accurately assess the variance over a region, due to the covariance among SNPs. To provide a neighborhood-level summary of SNP assignments to the four effect classes, we propose the following sliding-window statistic for SNP *i*, that we will call Weighted Cumulative Inclusion Probability (CIP_i_):

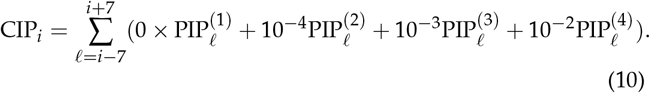

Finally, we used the Lewontin *D′* statistic (19) to quantify the LD between SNPs. Briefly, the LD coefficient *D_AB_* between SNPs A and B is defined as *D_AB_* = *p_AB_* – *p_A_p_B_*, where *p_A_*, *p_B_* and *p_AB_* respectively denote the frequency of allele A in the first locus, allele B in the second, and the frequency of simultaneously having both. *D*′ normalizes *D* so that 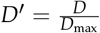, with

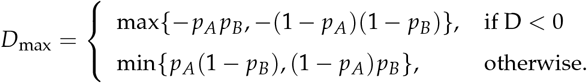

We will use the maximum value of the LD of a QTL with its neighboring SNPs as a reference for the link disequilibrium in the region.

##### Criteria ranking for QTL mapping

For the quantitative criteria *V_i_* and CIP*_i_* defined in Equations (9), (5) and (10), we propose the use of rankings for SNP prioritization rather than fixing value thresholds. For QTL mapping based the estimated posterior variance *V_i_*, we focus on the ten SNPs with the highest *V_i_*. As CIP*_i_* represents a sum over 15 SNPs in the neighborhood of SNP *i*, SNPs adjacent to those that are frequently categorized as non-null are likely to share large values for this criterion. As such, to address this redundancy, we focus on the 150 SNPs with the highest CIP*_i_* value.

### Data Availability

The Montbéliarde genotyping data on which simulations are based originate from a private French genomic selection program and were funded by the users (breeding companies and breeders). They are thus proprietary data that cannot be publicly disseminated to the scientific community. All code used to simulate and analyze the data, as well as the scripts to implement BayesR are available on GitHub (https://github.com/fmollandin/BayesR_Simulations).

**Table 2.**
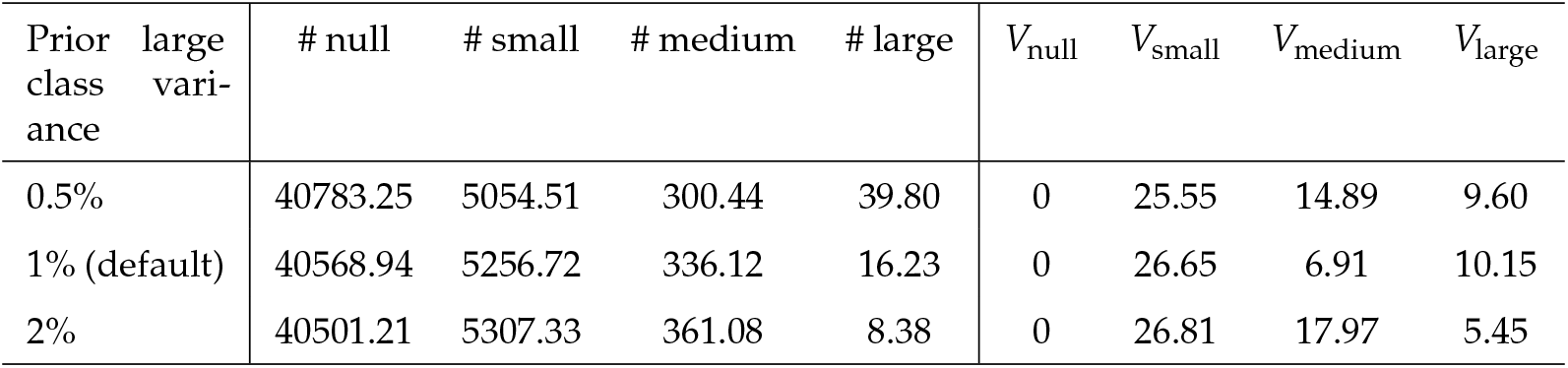
Average (across all simulation scenarios and independent datasets) of the posterior mean cardinality of each BayesR SNP effect class (null, small, medium, large) for three parameterizations of the prior large effect class variance. For a given dataset, each class size (#) is computed as the posterior mean of the number of SNPs assigned to each class across iterations, and *V_j_* is the posterior estimated cumulative variance of each class *j*.

| Prior large variance | # null | # small | # medium | # large | $V_{\text{null}}$ | $V_{\text{small}}$ | $V_{\text{medium}}$ | $V_{\text{large}}$ |
| --- | --- | --- | --- | --- | --- | --- | --- | --- |
| 0.5% | 40783.25 | 5054.51 | 300.44 | 39.80 | 0 | 25.55 | 14.89 | 9.60 |
| 1% (default) | 40568.94 | 5256.72 | 336.12 | 16.23 | 0 | 26.65 | 6.91 | 10.15 |
| 2% | 40501.21 | 5307.33 | 361.08 | 8.38 | 0 | 26.81 | 17.97 | 5.45 |

## RESULTS AND DISCUSSION

### Results

In the following, we first investigate the sensitivity of BayesR to parameter specification. We next evaluate the model’s performance for phenotype prediction and QTL mapping, based on the statistical criteria defined in the previous section, using simulated data that include a set of *n*_QTL_ = 5 QTLs, as well as polygenic SNPs and null SNPs with no effect on the phenotype.

#### Sensitivity of BayesR parameter specification

Although the proportion of additive genetic variance assigned to the small, medium, and large effect classes is typically set to 0.01%, 0.1%, and 1% respectively (see Equation (2) and (12)), these prior parameters can be varied by the user. To evaluate the impact on downstream results, we varied the latter between 0.5%, 1%, and 2% for all scenarios with *h*^2^ = 0.5, leaving those of the small and medium effect classes at their default values. Modifying the proportion of genetic variance of the large effect class did not appear to have a strong impact on the validation correlation; netherless we have observed differences in correlation among the three prior values that can reach 2.6% and 1% for the 50k and 50k custom data respectively. However, we do note that the posterior mean of the number of SNPs assigned in each class and its associated posterior estimated variance appear to be somewhat affected by this parameterization (Table 2). To assess the impact of the prior specification on per-SNP effect estimates, we calculated the Pearson correlation between the estimated posterior means 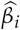 across SNPs, simulated scenarios and datasets. Among the three prior specifications, the correlation of estimated SNP effects was between 97.4% and 98.6% for all SNPs.

Based on these results, we consider that the prior specification appears to have little practical impact on the performance of BayesR, whether for its predictive performance or for per-SNP effect estimates. For the remainder, we therefore use the default prior specification for proportion of genetic variance in each effect class.

#### Predictive power of BayesR in varied simulation settings

We next sought to investigate the predictive power of BayesR across simulation scenarios, varying the contribution of QTLs to the additive genetic variance (which we refer to as scenarios below), heritability, and use of 50k or 50k custom genotype data.

The mean validation correlation (over the ten independent datasets simulated for each) for each simulation scenario illustrates the expected drop in prediction quality for decreasing heritabilities, whether 50k or 50k custom data are used (1). For the former, the mean (± sd) validation correlation across scenarios is 0.125 (±0.048), 0.301 (±0.057), 0.447 (±0.058) and 0.650 (±0.049) for *h*^2^ = {0.1,0.3,0.5,0.8}. For the latter, the inclusion of the true QTLs among the genotypes unsurprisingly leads to higher validation correlations, with mean (± sd) values across scenarios equal to 0.128 (±0.049), 0.312 (±0.058), 0.466 (±0.059) and 0.680 (±0.046) for *h*^2^ = {0.1,0.3,0.5,0.8}.

Although the trends of the mean validation correlation are non-linear as the QTL effects take on an increasing percentage of genetic variance for both types of data, we do remark an increasing disparity in performance between the 50k and 50k custom data, particularly as the heritability itself increases (2). In particular, as expected the potential gain in including the true causal mutations among genotypes (as is the case of the 50k custom data) appears to be particularly strong for moderate to large heritabilities and QTL effects. For *h*^2^=0.01, the average difference in validation correlation was 0.003 (±0.009), and in some cases the use of the 50k custom data actually corresponded to a slightly worse prediction. Similar results are observed at this level of heritability regardless of the simulated effect size of the QTLs. However, for *h*^2^ = {0.3,0.5,0.8}, 50k custom data led to a nearly systematic gain in performance: the average increase in validation correlation was 0.011 (±0.014), 0.019 (±0.020) and 0.031 (±0.030) across QTL effect size scenarios, and attained maximum values of 0.076, 0.112, and 0.160 respectively. For a given heritability, Figure 2 also shows marked improvements in prediction when including QTLs simulated with large shares of additive genetic variance.

**Figure 1.**
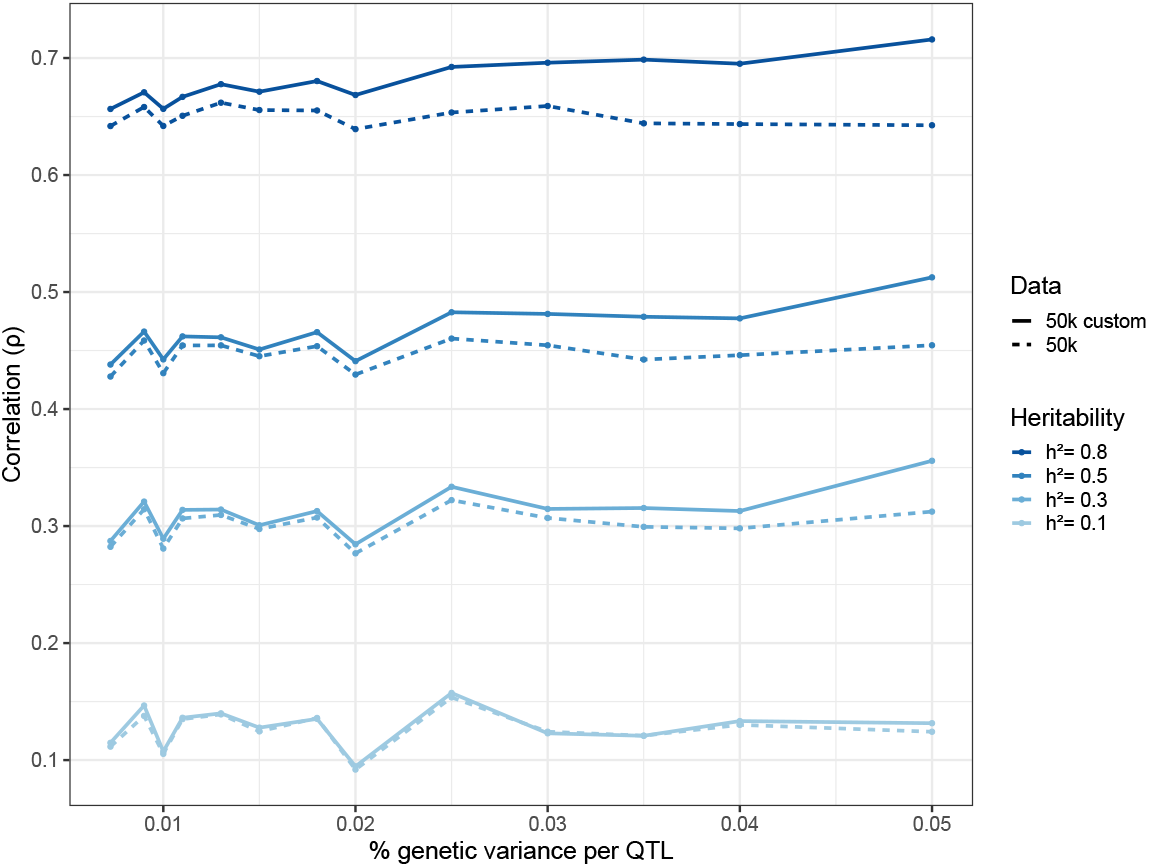
BayesR predictive performance across simulation settings. For each setting (*h*^2^ and percentage of genetic variance assigned to each QTL), points represent mean validation correlations across 10 independent datasets. Heritability values are represented by dark to light blue (*h*^2^ = 0.8 to 0.1), and solid and dotted lines represent results for the 50k and 50k custom datasets, respectively.

**Figure 2.**
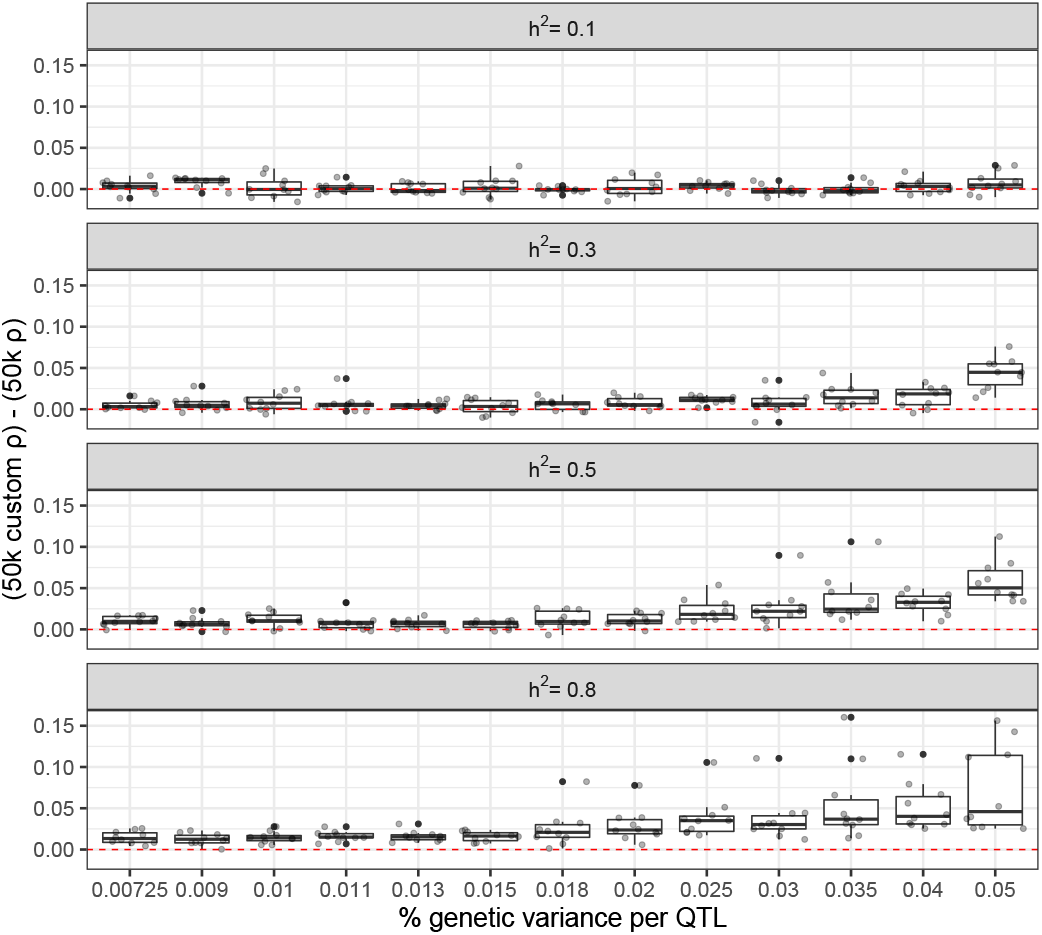
Difference in BayesR predictive performance for the 50k versus 50k custom genotypes across simulation settings. Each panel from top to bottom represents a given heritability (*h*^2^ = 0.1 to 0.8), and boxplots represent the distribution of differences in validation correlation between the 50k and 50k custom datasets for each independent dataset (i.e., for which the same 5 QTLs are simulated). The red dotted line indicates a baseline of 0.

#### QTL mapping using BayesR

A natural first tool to investigate for QTL mapping is the neighborhood PIP defined in Equation (6). We focus on the behavior of the neighborhood PIPs for the true QTLs across scenarios (3), averaging over the 50 QTLs available for each (5 QTLs × 10 independent datasets); note that as this is a windowbased measure, this measure can be computed for the true QTLs whether the 50k or 50k custom data are used. As shown in 3, the allotment of true QTL neighborhoods to effect classes varies widely across heritabilities, proportion of genetic variance for each QTL, and type of data used. Globally, assigning QTL neighborhoods to non-null effect classes, particularly the large effect class, is more frequent for larger heritabilities and simulated QTL effect sizes, as well as for 50k custom compared to 50k data. However, this difference disappears for small heritabilities; when *h*^2^ = 0.1, the average (± sd) neighborhood PIP for the null class across scenarios is 0.91 (±0.009) and 0.90 (±0.013) for the 50k and 50k custom data, respectively. Across scenarios, we observe a similar usage of the small effect class, with an average corresponding neighborhood PIP of 0.08 (±0.007) regardless of the genotyping data used. When *h*^2^ = {0.3,0.5,0.8}, as the simulated share of genetic variances for QTLs increases for both the 50k and 50k custom data, the null neighborhood PIP decreases and the large-effect neighborhood PIP increases. Across all simulated datasets and scenarios, the average (± sd) small- and medium-effect neighborhood PIPs are 0.117 (±0.053) and 0.058 (±0.040) respectively, illustrating that these two classes appear to be less often filled compared to the null and large classes (although all four classes do appear to be used outside of the lowest heritability setting).

The neighborhood PIP results provide a preview of how QTLs are grouped into non-null effect classes according to the neighborhood MAP rule (Equation (8); 4). In all simulation settings, no QTL neighborhoods were assigned to the small effect class using this criterion. When *h*^2^ = 0.1, without surprise, all QTLs were classified as null. For *h*^2^ = 0.5, a very small number of QTL neighborhoods were assigned to the medium effect class for the 50k data; increasing to *h*^2^ = 0.8 led to a larger number moving to this class for both the 50k and 50k custom data. When not assigned to the null class, it was much more common to attribute QTL neighborhoods to the large effect class; the number of correctly identified QTL neighborhoods increased with the simulated effect size and/or heritability, as well as when the causal markers were included among the genotypes; what’s more, these gains tend to accumulate when taken together. Correctly detecting at least one QTL window with the MAP rule required the proportion of genetic variance simulated for each QTL be using the 50k data, increasing to up to 6 QTL windows for larger simulated effects. A larger heritability of *h*^2^ = 0.5 for the same data required only *k* ≥ 0.9% to correctly identify at least one QTL window, which increases to 22 for *k* = 5%. However, including the causal markers in the genotype data enabled detection of QTL windows at *k* ≥ 1.3% for *h*^2^ = 0.3, with up to 30 correctly detected at *k* = 5%. In the most favorable scenario, with *h*^2^ = 0.8 and 50k custom data, QTL windows are detected for all values k, and they are exhaustively assigned to the large effect class for *k* = 5%.

Given these results, it is not surprising that the neighborhood *MAP*^non–nuLL^ in Equation (8) will tend to detect more QTL windows as being non-null. However, it is also useful to consider the behavior of this criterion while considering the LD blocks specific to each simulated QTL. In 5, we visualize the neighborhood inclusion probability IP*_i_* (defined in Equation (7)) for each of the 50 simulated QTL windows across scenarios for *h*^2^ = 0.5, illustrating the proportion that are correctly included as non-null in the model (i.e., when the neighborhood inclusion probability > 0.5). The *MAP*^non–null^ appears to require a minimum LD of 55% to correctly recover QTL windows using the 50k data. Below this threshold, a large portion of QTL windows are not detected. Above this threshold, QTL window detection appears to become feasible once the simulated per-QTL percentage of genetic variance attains about *k* = 2%. In the 50k custom data, QTL window detection does not however depend on the amount of LD, although we do note lower inclusion probabilities for QTLs in very high LD with their neighbors as compared to the 50k data. Similar to the 50k data, there is an effect size threshold at about *k* = 1.8% at which QTL windows are more frequently detected.

Because the same five QTLs are simulated in each independent dataset across effect size scenarios, Figure 5 also allows for their specific detection to be followed across configurations. Thus, it can be seen that some QTLs windows are not detected in any of the scenarios, while others are more easily detected, even for lower shares of the genetic variance. That said, there are occasionally discontinuities in detection observed for increasing shares of the variance (i.e., a QTL window correctly identified for *k* = 0.02 but not 0.025). With the exception of *h*^2^ = 0.1, which had very weak detection in all scenarios and datasets, we found similar conclusions for *h*^2^ = 0.3 and 0.8, with respectively slightly smaller and larger overall inclusion probabilities than those shown in Figure 5.

**Figure 3.**
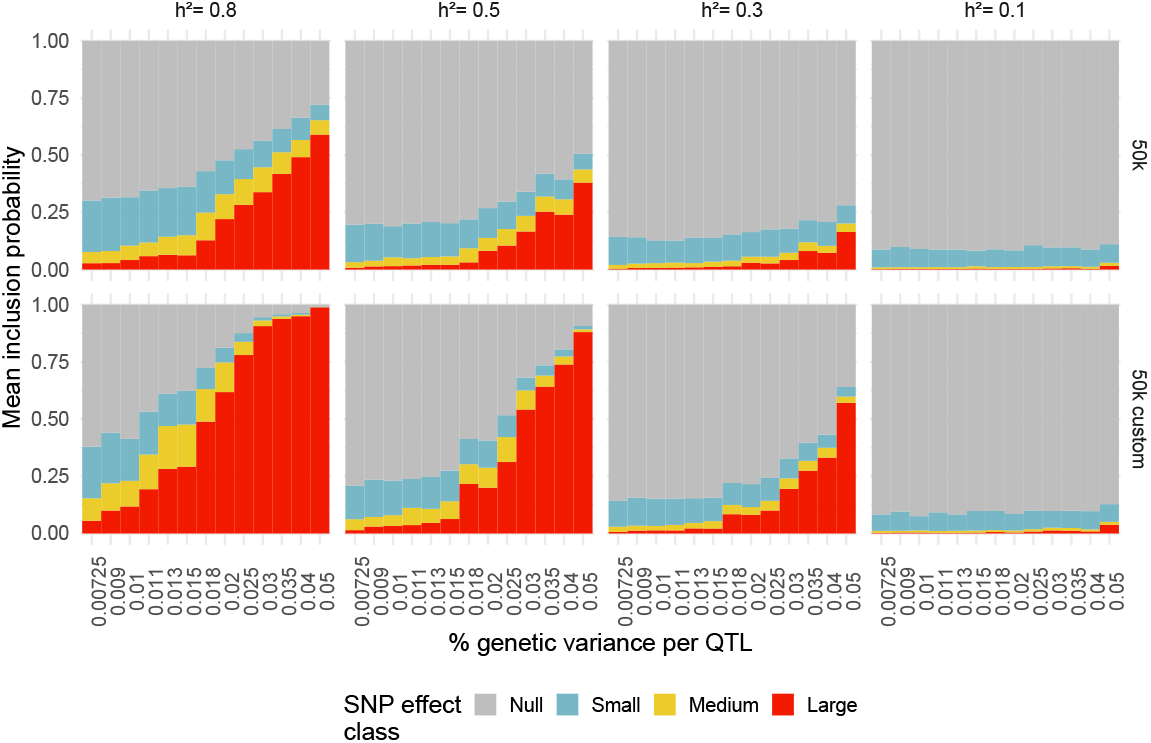
Neighborhood posterior inclusion probabilities across simulation settings. Panels represent combinations of heritability (columns; *h*^2^ = 0.8 to 0.1) and type of data used (rows; 50k or 50k custom). Bars represent average (across 5 QTLs × 10 independent datasets) neighborhood PIP values for the four BayesR effect size classes: null (grey), small (blue), medium (yellow), and large (red).

**Figure 4.**
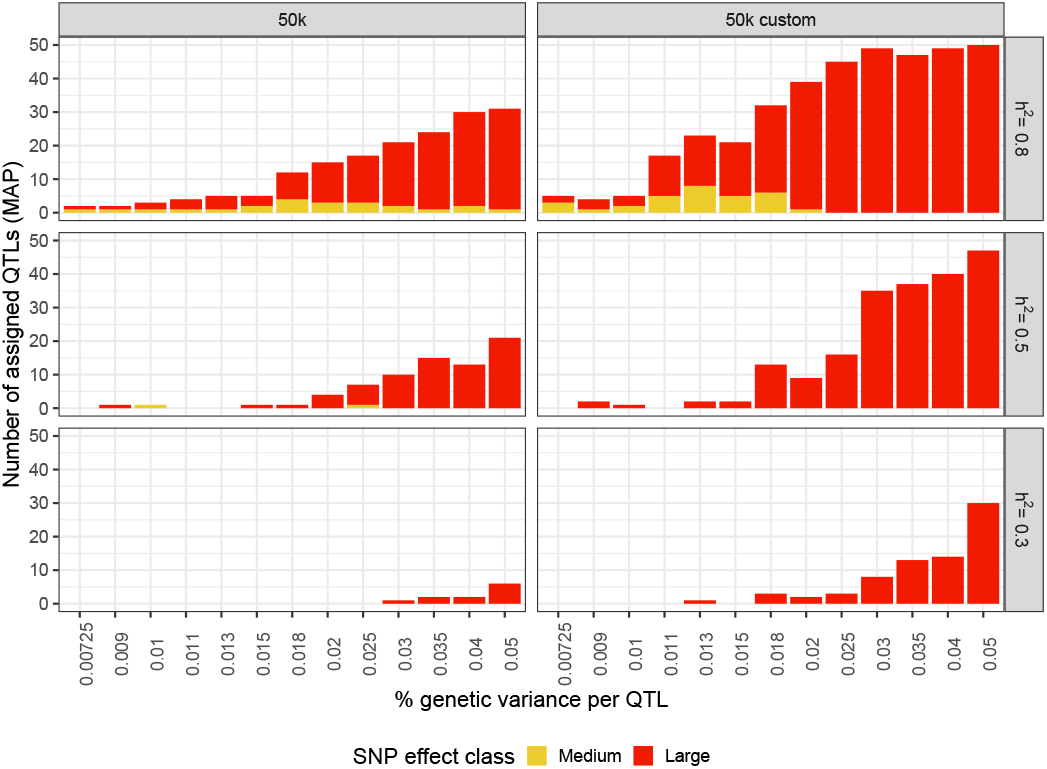
Neighborhood MAP rule for QTL mapping across simulation settings. Number of true QTL windows (out of 5 QTLs × 10 independent datasets simulated for each scenario, corresponding to a total of 50) correctly assigned to the medium (yellow) and large (red) effect size class using the neighborhood MAP rule. Panels represent data type (columns; 50k and 50k custom) and heritability (rows; *h*^2^ = 0.8 to 0.1). The small effect class is not represented because it was empty across all simulation configurations.

**Figure 5.**
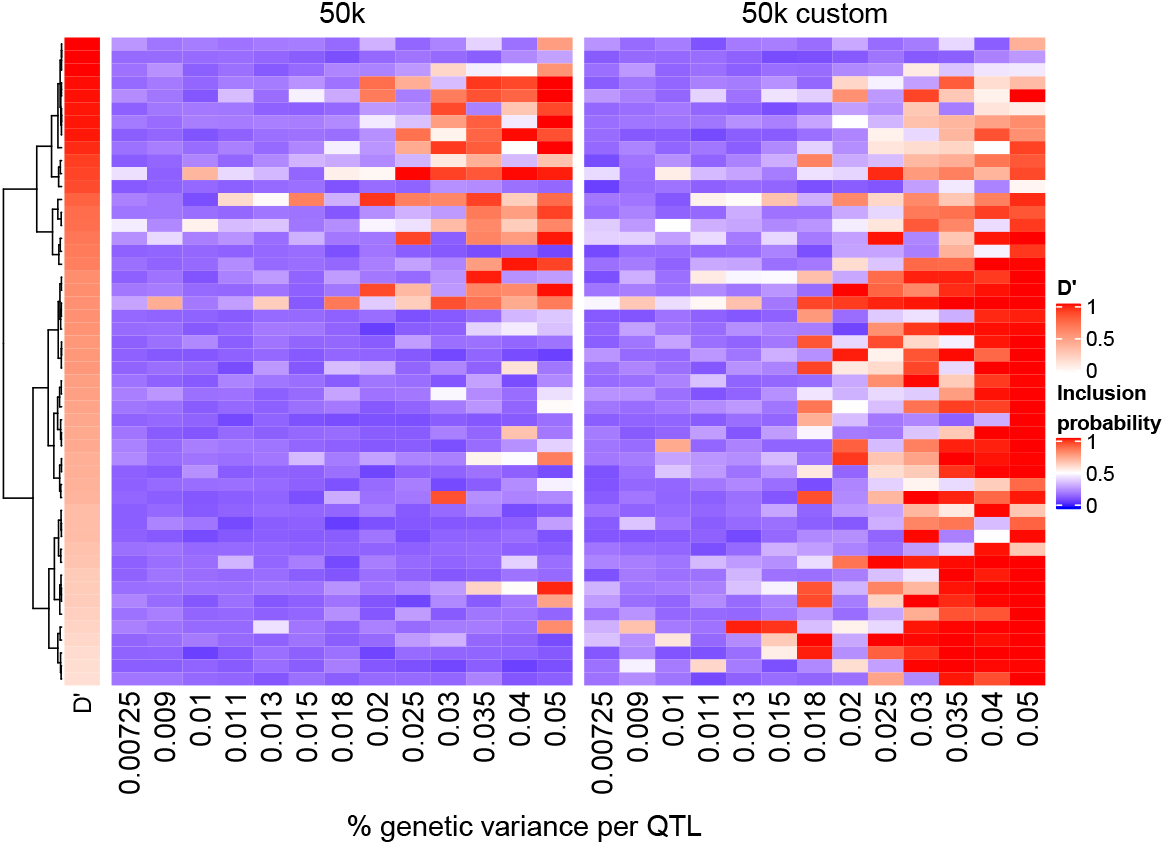
QTL window mapping using the neighborhood inclusion probability across different effect sizes and LD strengths for *h*^2^ = 0.5. Neighborhood inclusion probabilities 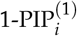 for each of the 50 simulated QTLs (heatmap rows) for the 50k (left) and 50k custom (right) data across scenarios (heatmap columns). QTLs are sorted in descending order according to their LD, as measured by *D*′ (left annotation, with deeper reds representing larger values). QTL windows that are represented by white to red cells are correctly detected using the neighborhood non-null MAP.

**Figure 6.**
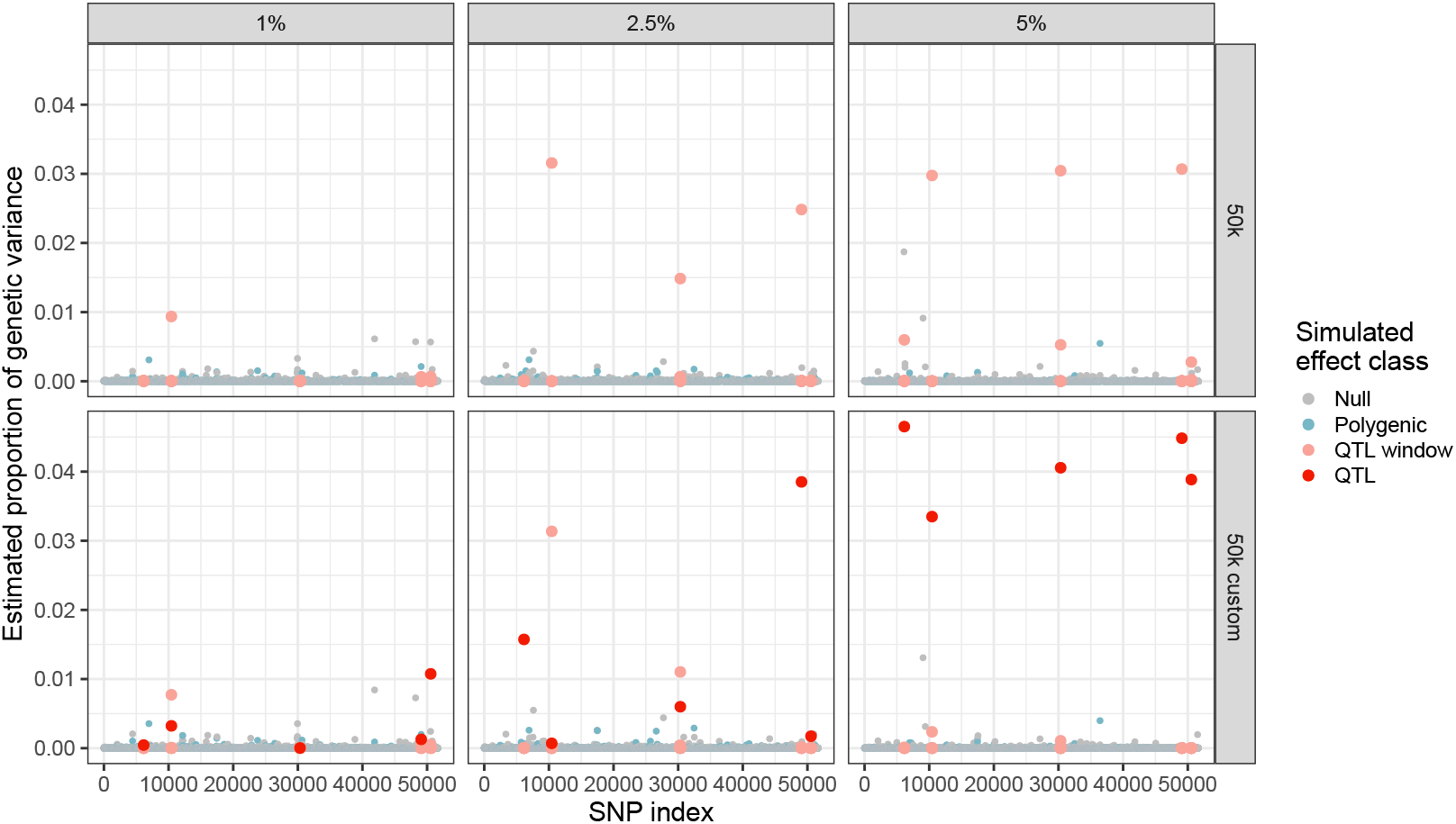
Genome-wide posterior estimate of the proportion of genetic variance per SNP for a single dataset with *h*^2^ = 0.5. Posterior estimates of the per-SNP proportion of genetic variance across all *p* = 46,178 SNPs for one of the simulated independent datasets. Panels represent a given simulation setting for percentage of genetic variance per QTL (columns; *k* = {1%, 2.5%, 5%}) and data type (rows; 50k versus 50k custom). Points represent individual SNPs, and are colored according to their true effect class (null, polygenic, in the neighborhood of a true QTL, and true QTL). The same five QTLs appear in each panel; true QTLs are only present in the 50k custom data.

**Figure 7.**
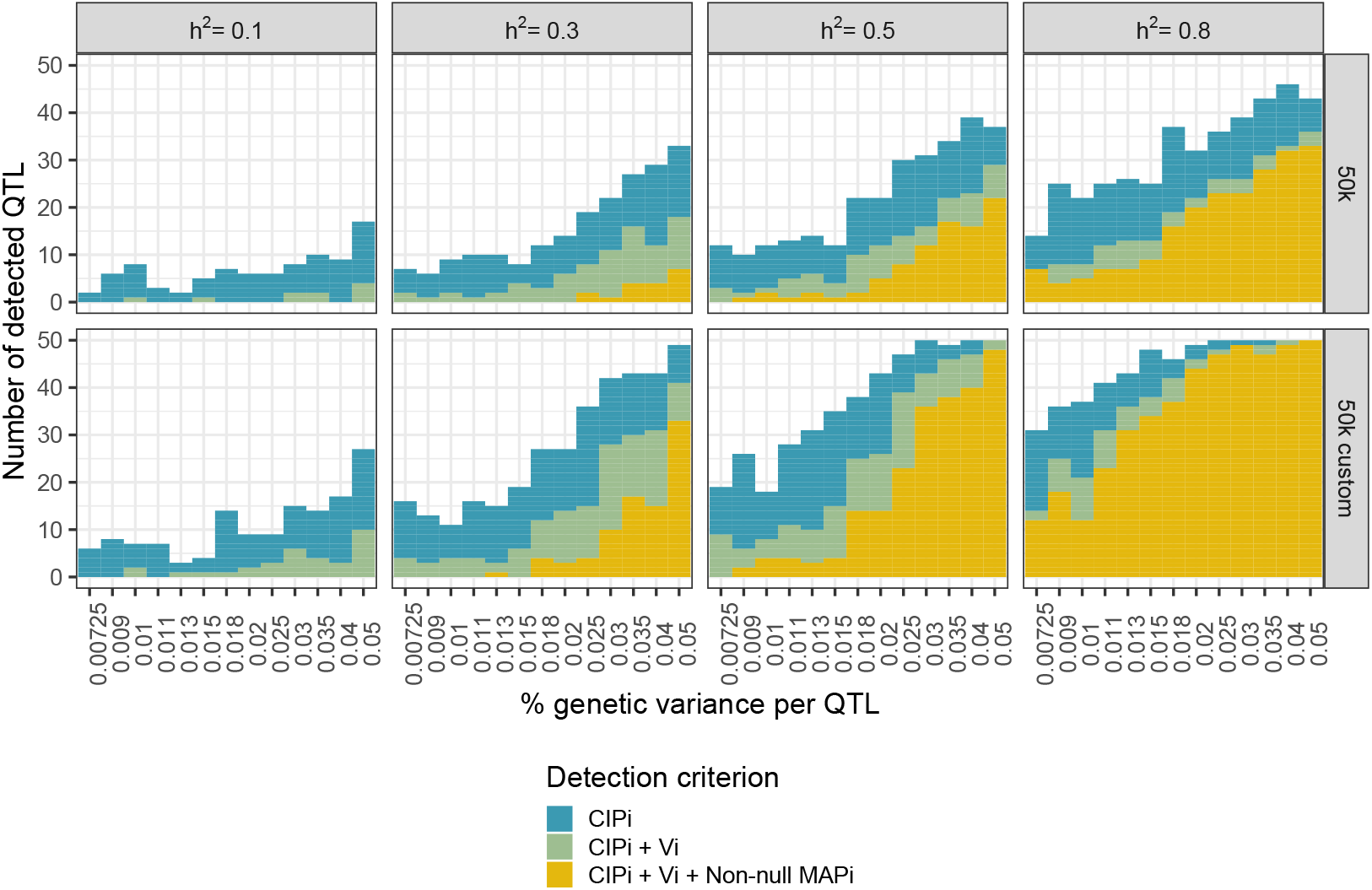
QTL window mapping using three different criteria across simulation settings. Number of true QTL windows (out of 5 QTLs × 10 independent datasets simulated for each scenario, corresponding to a total of 50) corrected identified using the CIPi ranking (top 150), *V_i_* (top 10), and 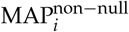 neighborhood criteria. Panels represent data type (rows; 50k and 50k custom) and heritability (columns; *h*^2^ = 0.1 to 0.8).

Beyond the assignment of SNPs to effect classes using the neighborhood PIPs (and corresponding MAP rules), BayesR also provides posterior estimates of variability at several levels, including the additive genetic variance 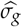, the cumulative variance for each of the three non-null effect classes, and the variance of each SNP. Before discussing the latter (arguably the most pertinent for QTL mapping), we verify the estimation quality of the additive genetic variance. In the 50k genotype data, on average (± sd) across scenarios, 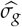 was 9.06 (±3.32), 30.85 (±3.93), 50.12 (±4.30) and 77.36 (±4.61) for *h*^2^ = {0.1,0.3,0.5,0.8} respectively; the corresponding true value of *σ_g_* for each were 10, 30, 50 and 80. In the case of the 50k custom data, this same parameter was estimated to be 9.11 (±3.27), 31.01 (±3.97), 50.27 (±4.32) and 77.54 (± 4.49), respectively.

Given that the total additive genetic variance appears to be well-estimated for both types of genotype type, we turn our attention to the posterior variance 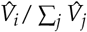 of each neighborhood as defined in Equation (9). We focus in particular on the case where *h*^2^ = 0.5 and proportions of genetic variance per QTL equal to *k* = {1%,2.5%,5%} (6); similar trends were observed for *h*^2^ = {0.3,0.8}. We note that the estimated proportion of genetic variance per SNP window are largely shrunk towards zero, clearly distinguishing those included in the model. In the 50k data, certain true QTL windows are clearly prioritized and easily identifiable. Of the 5 simulated QTLs, we observe one that can be visually identified for *k* = 1%, and three for *k* = {2.5%,5%}; more moderated peaks are observed for the remaining QTLs. In addition, the estimated posterior SNP window variance is about 3%, regardless of the share of variance for the simulated QTLs. When *k* = {1%, 2.5%}, the prioritized QTL windows appear to have estimated variances close to the true simulated values. These estimates further improve when the 50k custom data are used, and a larger number of QTLs are clearly prioritized: we note that 2, 4 and 5 QTLs have visibly distinct peaks for *k* = {1%,2.5%,5%}, respectively.

As a final criterion, we investigate the weighted cumulative inclusion probability statistic CIP*_i_* defined in Equation (10) as a way to prioritize neighborhoods where the assignment of SNPs to non-null classes is somewhat diluted. This statistic tends to up-weight regions as SNPs in the neighborhood are assigned to non-null classes (potentially in the place of the primary QTL, which may be in tight LD with its neighbors). We expect QTL windows already detected by the neighborhood MAP to similarly have large CIP*_i_* values; however, it may facilitate the detection of those for which a cumulative integration of non-null SNPs across the window provides additional information.

To evaluate this point, we compared the QTL mapping performance of BayesR using the following three criteria: the neighborhood 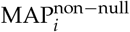, and the rankings of the neighborhood *V_i_* (top ten) and neighborhood CIP*_i_* (top 150). We chose to use 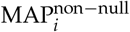 here rather than MAP*_i_* as it is less stringent. Across simulation scenarios and heritabilities, all QTL windows correctly detected by the non-null neighborhood MAP were also identified by the other two criteria (7). Similarly, all QTL windows correctly detected by the posterior neighborhood variance V*_i_* ranking were all also flagged by the CIP*_i_* ranking. The sliding window statistic thus appears to provide the greatest detection sensitivity, while the MAP criterion is the most conservative.

For all three criteria, the number of detected QTLs increases with the simulated effect size and heritability, as well as with their inclusion among the genotypes (50k custom data), with the exception of the lowest considered heritability, *h*^2^ = 0.1. In this case, no QTL windows are detected with the MAP^non–null^, and the number of QTLs identified does not greatly increase for larger QTL effect sizes. Using the CIP*_i_* rankings, about half of the true QTL windows can be recovered using the 50k data when *h*^2^ = 0.8 in the 50k chip, and similar results are possible with the 50k custom data for *h*^2^ = 0.5. When the true QTLs are excluded from the genotypes, at most 46 of the 50 true QTL windows can be identified with CIP*_i_*, even in ideal circumstances (h^2^ = 0.8 and *k* = 4%). However, using the 50k custom data that include these QTLs allows for universal detection when *h*^2^ = 0.5 and *k* = {3%, 4%, 5%}, or *h*^2^ = 0.8 for *k* ≥ 2.5%.

## Discussion

In this work, we evaluated the performance of the BayesR Bayesian genomic prediction model for prediction quality and QTL mapping performance on simulated data under a variety of scenarios, including varying QTL effect sizes, heritabilities, and the use of 50k versus 50k custom genotype data. Simulated phenotypes were generated using SNPs from a real set of genotype data in cattle that were divided into three categories (null, polygenic SNPs, and QTLs), with variable corresponding shares of the additive genetic variance. In our study, polygenic SNPs were simulated to have the same share of genetic additive variance as the default BayesR small effect class, i.e. 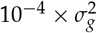. QTLs were assigned variances ranging from 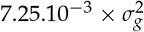 to 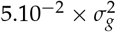, constituting an interval that includes the default prior variance of the BayesR large effect class, i.e. 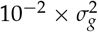. These scenarios were simulated at different levels of heritability *h*^2^ = {0.1,0.3,0.5,0.8}, and we considered both genotype data that excluded (50k data) or included (50k custom data) the true simulated QTLs. As the BayesR model definition includes four different effect size classes (null, small, medium, and large), it is of particular interest to evaluate how well the model itself adapts to the underlying genomic architecture of the data.

The specific parameterization of BayesR (e.g., number and magnitude of non-null effect classes) can be adapted for different applications. In this work, we investigated the sensitivity of BayesR results based on the magnitude of the large effect class, and we found that the performance of BayesR (predictive power, estimations of per-SNP effects) was relatively robust. This suggests a limited benefit to modifying the priors based on prior biological knowledge. A more promising approach to integrate such prior knowledge is the related BayesRC model (13). In the BayesRC approach, SNPs are divided by the user into two or more non-overlapping subsets, each of which represents a biologically relevant grouping with a potentially different proportion of QTLs. For each subset, the four BayesR SNP effect classes are used, with proportions modeled using an independent Dirichlet prior (i.e., varying among subsets). As this flexibility can help prioritize informative SNP subsets that contain a larger proportion of QTLs, it would be of great interest to evaluate the impact of the choice of SNP subsets on QTL mapping with BayesRC, using the criteria we investigated here.

With the exception of very low heritability (*h*^2^ = 0.1), validation correlation unsurprisingly increases when QTLs are included among the genotypes (i.e., the 50k custom data); this increase is particularly marked for highly heritable phenotypes as well as for QTLs with large effects. We note that the predictive power of the BayesR model varied both across simulated scenarios, as well as within a given scenario, suggesting that the specific position of simulated QTLs and polygenic SNPs appears to have an influence on the behavior of BayesR.

We presented several statistics for QTL mapping and interpretation using BayesR results, but we note that accurately assessing and quantifying the importance of a particular genomic region remains a challenge. One major obstacle is the presence of LD between SNPs. On one extreme, low LD among neighboring SNPs can impede the detection of regions if causal mutations are not directly included among genotypes, while on the other, strong LD blocks can dilute the signal among adjacent SNPs, leading to alternating assignments to non-zero effect classes (and subsequently lower estimated PIPs and variances). While the 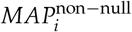 appears to be overly conservative for the detection of QTL neighborhoods, the *V_i_* has the advantage of facilitating an estimation of the proportion of variability corresponding to each QTL neighborhood, given the overall estimated genetic additive variability. On the other hand, the *CIP_i_* statistic better takes LD into account by incorporating the cumulative importance of an entire region, perhaps explaining why it can better identify QTL neighborhoods than the other criteria considered, even under non-optimal conditions (e.g. *h*^2^ = 0.1).

There are several limits to our current study that should be taken into consideration. First, we note that some of our simulation scenarios could be considered to represent optimal conditions (e.g., large heritabilities and QTL effect sizes) that would be rare in real applications. However, studying these extreme scenarios enables the behavior of the BayesR model to be established in ideal cases. All of our simulations made use of a constant number of individuals in both the training and validation sets, but a future study evaluating the impact of the training population sample size on QTL mapping ability, particularly for cases with low heritability (e.g. *h*^2^ = 0.1), could provide insight on this point. Lastly, when sampling SNPs to represent QTLs in our simulations, we chose to limit the choice to those with a MAF > 0.15, thus excluding those with rare alleles. Although this allowed us to avoid edge cases that would arise with very low MAFs, making it easier to homogenize simulated datasets across different selections of QTLs, this however is an important consideration in QTL mapping.

## CONCLUSION

BayesR is a powerful tool for simultaneously providing accurate phenotypic predictions and mapping causal regions. Our simulation results illustrate the flexibility of BayesR for different genomic architectures for all but very low heritabilities (*h*^2^ = 0.1) or small QTL effects (<1% share of the additive genomic variance). Although the four effect size classes (null, small, medium, large) defined in BayesR do not themselves always reflect the true categorization of SNPs, they do offer a new approach to understanding and characterizing the genomic architecture underlying a phenotype. To this end, we presented a variety of statistical criteria that can be used to perform QTL mapping using the output of the BayesR model, including neighborhood-based non-null maximum a posteriori rules, posterior estimated variances, and cumulative inclusion probabilities. We showed that some of the challenges in QTL mapping posed by strong LD blocks could be overcome using the latter criterion, which focuses on the assignment to non-null effect classes of SNPs in an entire neighborhood. By ranking SNPs using this criterion, we demonstrated that QTL windows could more easily be detected, even in simulation scenarios with more challenging conditions.

## Acknowledgements

The authors thank Mario Calus, Marco Bink, and Bruno Perez for helpful discussions, and Didier Boichard for providing the simulation software used to generate simulated phenotypes from a set of genotype data.

## ADDITIONAL FILES

All code used to simulate and analyze the data, as well as the scripts to implement BayesR are available on GitHub (https://github.com/fmollandin/BayesR_Simulations). The repository is divided into three parts:

- **Simulations** Fortran source code of the software used to simulate data based on real genotypes. An example of parameters and description of their use are also provided.
- **bayesR** The modified version of BayesR (available at https://github.com/syntheke/bayesR), including recovery of the estimated per-SNP effects for each iteration, which in turn facilitates the estimation of per-SNP posterior variances. An example of the use of this software is also provided.
- **codes_R** Partial R scripts used to analyze the BayesR model output and visualize the corresponding results. Scripts to reproduce all figures presented in the article are also included.

